# Peptide generative design with weakly order-dependent autoregressive language model and lifelong learning

**DOI:** 10.1101/2023.12.31.573750

**Authors:** Zhiwei Nie, Daixi Li, Yutian Liu, Fan Xu, Hongyu Zhang, Xiansong Huang, Xudong Liu, Zhennan Wang, Yiming Ma, Yuxin Ye, Feng Yin, Wen-Bin Zhang, Zhixiang Ren, Zhihong Liu, Zigang Li, Jie Chen

## Abstract

Bioactive peptides have become strong candidates for a variety of clinical therapies due to their diverse advantages, which promotes the development of deep generative models for peptide design. Considering that existing methods cannot effectively deal with the conformational flexibility of peptides and are difficult to capture accurate residue-to-residue interaction dependencies, we propose a unified weakly order-dependent autoregressive language modeling architecture (PepGenWOA) for bioactive peptide generative design with tolerating out-of-order input as inductive bias. The superiority of PepGenWOA is demonstrated by generating three classes of therapeutic peptides, including antimicrobial peptides, anticancer peptides, and peptide binders. For antimicrobial and anticancer peptide generation, PepGenWOA not only comprehensively outperforms state-of-the-art baseline models, but also exhibits a significant propensity to incorporate specific types of residues that are beneficial for antimicrobial or anticancer bioactivity. For characteristic-guided peptide binder generation, the pretrained PepGenWOA is fine-tuned with Mixture-of-Experts-style plugins with lifelong learning paradigm, thereby achieving the best trade-off between memory stability for world knowledge and learning plasticity for new downstream tasks. The subsequent structure-based virtual screening anchors the 7 most promising candidates from the mega-scale synthetic sequence space, reducing the number of candidates for *in vitro* experimental validation by 5 orders of magnitude. A target binding rate of 28.6% with binding specificity further confirms the effectiveness of the fine-tuning strategy. Overall, PepGenWOA is a unified architecture for peptide generative design that can be flexibly customized for different task requirements, thereby harnessing generative language models to reach the “dark space” of biological language.

## 1 Introduction

Bioactive peptides play a wide range of roles in human physiology [1, 2], acting as hormones, neurotransmitters, growth factors, ion channel ligands, etc. [3–7] Compared with traditional small molecules [8], they have become an ideal starting point for the development of a variety of novel therapies due to their excellent safety, tolerability, and low production costs [9]. As an important category of active peptides, therapeutic peptides represented by antimicrobial peptides (AMPs) [10], anticancer peptides (ACPs) [11] and peptide-based binders [12], have shown great potential in dealing with complex human diseases. Traditionally, bioactive peptide design relies on rational design methods [13] for optimization, or employs computational tools, especially machine learning models, to mine candidates in the global microbiome [14, 15]. However, the potential peptide space is vast, making it difficult for the above approaches to efficiently anchor therapeutic sub-spaces in it. In addition, rational design methods and screening strategies cannot avoid significant costs in time and money.

With the booming development of deep generative models in fields such as vision [16, 17] and natural language [18, 19], they have also become popular in the field of protein design [20], especially bioactive peptide design [21]. From the perspective of generation mode, we can divide the existing peptide generative design studies into three categories: sequence-based generation, structure-based generation, and sequence-structure co-design. First, for peptide sequence generation, the autoregressive models [22–24] treat the peptide sequence as a sentence with residues as tokens, and generate residues one by one depending on previously generated residue arrangements. The sequence generation methods based on Variational Autoencoder (VAE) [25–28] employ an encoder-decoder architecture to learn the latent space and sample from it to generate peptide sequences, and conditional constraints can be added. The sequence generation methods based on Generative Adversarial Network (GAN) [29–31] train the generator and discriminator to compete with each other to achieve the generation of peptide sequences. The diffusion-model-based sequence generation methods [32, 33] estimate an unknown data distribution through a forward diffusion process and a reverse denoising process, and then generate peptide sequences through different samplers. Second, for peptide structure generation, many previous studies of protein structure generation can be naturally transferred and applied. For example, FoldingDiff [34], which employs the inter-residue angles in protein backbones, and ProtDiff [35], which generates protein backbone structures, are both diffusion-based generation models that can be extended to peptide structure generation. Third, for the sequence-structure co-design of peptides, a multi-modal diffusion model relying on contrastive learning [36] is proposed to generate AMPs and ACPs. In addition, the sequence-structure co-design methods for general proteins [37] or antibodies [38] can also be extended to peptide design.

However, existing generative models for bioactive peptide design have significant shortcomings. First of all, peptides are often partially or even completely unstructured in isolation [39, 40]. When the target structure is unclear or the binding mode is uncertain, the flexibility and instability of the peptide conformations [41, 42] are difficult to deal with by previous structure-based methods. In this case, the sequence-based generation model is more likely to internalize the dynamic information hidden in the sequences and implicitly capture the potential interaction mode with the target in the disordered peptide conformation ensembles. However, previous sequence-based generative methods treat all one-dimensional protein sequences as sentences of sequentially arranged residues and ignore the spatial proximity between residues caused by protein folding [43–45], which makes it difficult for the neural network to capture the correct residue-to-residue interaction dependencies. Specifically, as shown in Fig.1a, residues that are far apart in a one-dimensional protein sequence may become spatially close after folding, giving rise to diverse residue-to-residue interactions [46, 47], including hydrogen bonds, hydrophobic interactions, electrostatic interactions, and disulfide bonds, etc. Naturally, the above phenomenon has a more significant impact on the intramolecular interaction network of peptides with great conformational flexibility [41, 42]. In addition, diverse residue-to-residue interactions constitute different intramolecular interaction networks that share the basic mechanisms that support the peptide bioactivity execution (Fig.1b), which further emphasizes the necessity of capturing correct residue-to-residue interaction dependencies for peptide generative design. Therefore, generative language modeling methods for peptide design that learn and build correct residue-to-residue interaction dependencies from massive peptide sequences are urgently needed.

**Fig. 1.**
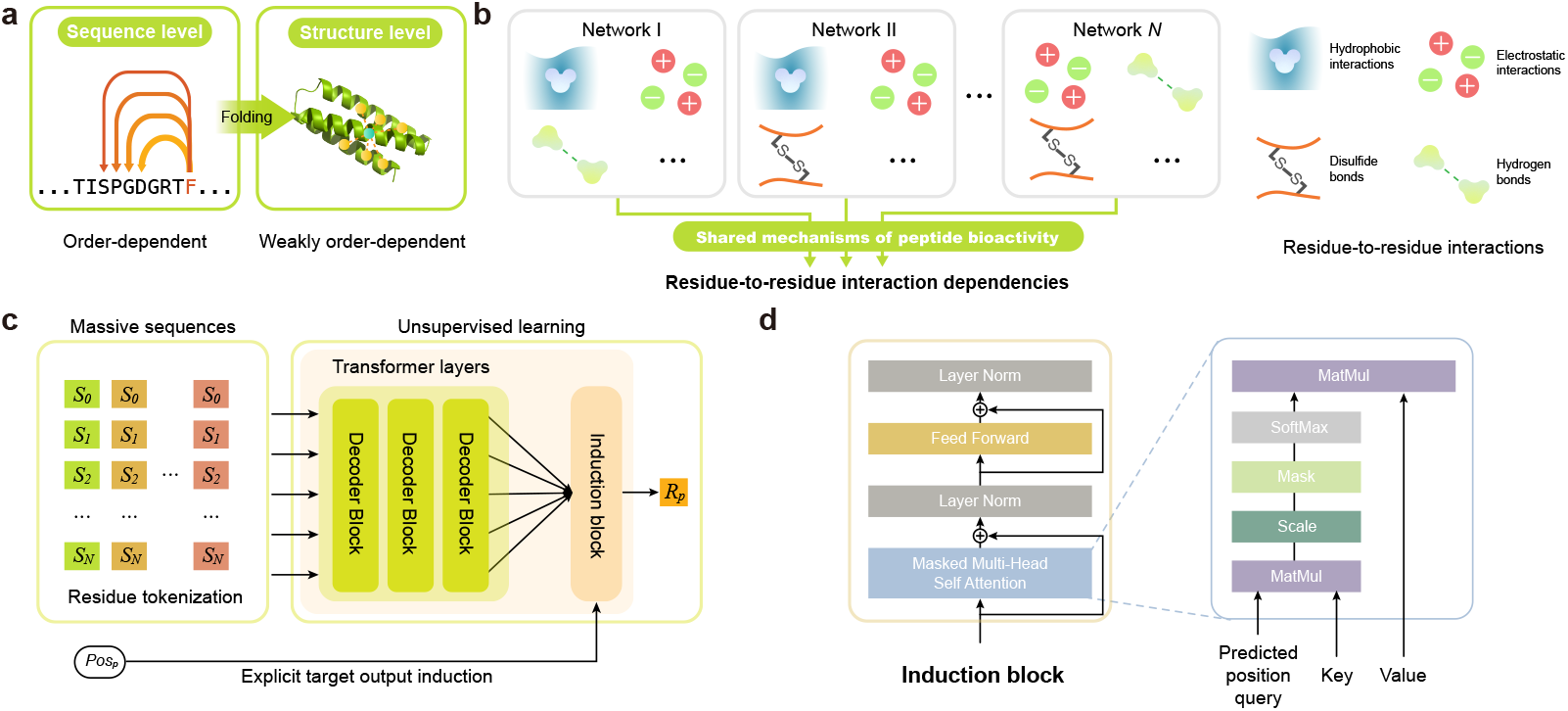
Motivation and model architecture of PepGenWOA. **a**, The necessity of capturing residue-to-residue interaction dependencies from one-dimensional protein sequences. Residues that are not adjacent in one-dimensional protein sequence may be spatially close to each other due to protein folding, but previous protein sequence generative modeling architectures ignored this phenomenon and treated the sequence as a sentence containing sequentially arranged residues, making it difficult for the neural network to learn the correct residue-to-residue interaction dependencies. **b**, Illustration of different intramolecular interaction networks, which consist of diverse inter-residue interactions, including hydrogen bonds, hydrophobic interactions, electrostatic interactions, and disulfide bonds. For bioactive peptides, different intramolecular interaction networks share the basic mechanisms of peptide bioactivity. **cd**, The weakly order-dependent autoregressive architecture of PepGenWOA, in which an induction block is designed to provide explicit target output induction, i.e., the position of predicted residue (c). *P os*_*P*_ represents the position of the predicted residue, and *R*_*P*_ represents the predicted residue. The induction block is a variant of the transformer decoder, in which an additional embedding indicting the next predicted position is employed as the query vector of the attention mechanism (d). The introduction of induction block allows the autoregressive model to be weakly order-dependent to tolerate out-of-order inputs, which enables the model to not only predict the next residue when training, but also understand the correct “semantics” of the random residue order (i.e., the correct interaction dependencies between residues that determine peptide bioactivity).

To this end, we propose a **W**eakly **O**rder-dependent **A**utoregressive language modeling architecture for bioactive **Pep**tide **Gen**erative design, abbreviated as Pep-GenWOA. PepGenWOA is capable of the materializing intramolecular interaction networks to identify their commonalities for bioactivity execution through unsupervised learning with tolerating out-of-order input as inductive bias. We demonstrate the superiority of PepGenWOA by generating three classes of therapeutic peptides, including antimicrobial peptides, anticancer peptides, and peptide binders. For the generation of antimicrobial and anticancer peptides, PepGenWOA comprehensively outperforms state-of-the-art baseline models. More importantly, PepGenWOA exhibits a significant propensity to incorporate specific types of residues that are beneficial for antimicrobial or anticancer bioactivity, demonstrating a level of internalizing the shared basic bioactivity mechanisms of therapeutic peptides. Furthermore, we design a characteristic-guided peptide binder generation pipeline with lifelong learning, in which five fine-tuning strategies are comprehensively evaluated to determine the best trade-off between memory stability for world knowledge and learning plasticity for new downstream tasks. Through fine-tuning the pretrained PepGenWOA with Mixture-of-Experts-style plugins, more than 2.2 million peptide binder candidates are generated and performed mega-scale structure-based virtual screening to efficiently anchor the 7 most promising candidates. *In vitro* experimental validations of these candidates show that the target binding rate is close to 29% with binding specificity. Overall, PepGenWOA represents a unified language modeling architecture for bioactive peptide generative design with broad applicability and customization flexibility.

## 2 Results

### 2.1 Weakly order-dependent autoregressive language modeling architecture

The autoregressive language modeling architecture is suitable for predicting the next residue at the sequence level, which is composed of a stack of transformer decoders [48–50]. In general, the autoregressive language model for peptide sequence generation directly outputs the residue type at a specific position after its previous time step, which leads to the prediction of the residue type at this position being strongly dependent on its previous residue order [49, 50]. This strong dependence on residue order is expected to severely weaken the ability of this architecture to capture correct residue-to-residue interaction dependencies through training on a large number of protein sequences. As shown in Fig.1c, we propose a weakly order-dependent autore-gressive language model, named PepGenWOA, for bioactive peptide generative design. The core difference between PepGenWOA and the original autoregressive language modeling architectures is the introduction of induction block, a variant of the transformer decoder. Take independent position encoding of the next predicted residue as induction, PepGenWOA not only predicts the next residue during training on massive protein sequences, but also understands the correct semantics of the random residue order, i.e., the correct interaction dependencies between residues that deter mine the peptide bioactivity. Specifically, the induction block is located above the stacked transformer decoder blocks and employs an additional next position embedding as the query vector of the masked multi-head attention (Fig.1d). The model therefore clearly knows the residue position to be predicted, allowing itself to capture the correct interaction dependencies between residues even when the input is out of order. By introducing tolerance for out-of-order inputs as an inductive bias of the autoregressive architecture, PepGenWOA understands and builds the intricate residue-to-residue interactions through unsupervised learning, thereby internalizing the shared basic mechanisms of the bioactivity execution of peptides. Therefore, Pep-GenWOA is expected to achieve the generation of specific types of bioactive peptides through pretraining and subsequent fine-tuning on different sets of peptide sequences.

### 2.2 Generation performance for AMP and ACP

Antimicrobial peptides (AMPs) and anticancer peptides (ACPs) are representative therapeutic peptides [1, 2]. As good alternatives to current antimicrobial drugs, AMPs have a broad range of target organisms [51, 52]. With significant antimicrobial activity and unique antimicrobial mechanism, AMP is the subject of continuous attention and research [51, 52]. In recent years, bioactive peptides with anticancer properties have emerged as a competitive therapeutic option because conventional therapies used to treat cancer produce significant side effects [53, 54]. Therefore, considering the positive significance of AMP and ACP for clinical treatment, we adopt these two types of therapeutic peptides to demonstrate the generation performance of PepGenWOA.

First, we implement various baseline models, including sequence-based generative models [22, 27–29] and sequence-structure co-design generative models [36–38], for the generation of AMPs and ACPs. For the generated AMP and ACP sequences, we adopt the alignment score [55] that reflects the similarity between generated sequences and the training set to evaluate their novelty. The smaller the alignment score, the higher the novelty of the generated sequence. Considering the unstable nature of peptide conformation, we adopt the instability score [56, 57] to evaluate the stability of the generated sequences. The smaller the instability score, the higher the stability of the generated sequence. Furthermore, to evaluate the therapeutic properties of the generated sequences, we adopt antimicrobial probability for AMP and anticancer probability for ACP respectively. Specifically, for the generated AMP sequences, we employ a binary classification model [58] to predict the probability that they are antimicrobial peptides. Similarly, for the generated ACP sequences, a binary classification model [59] is adopted to predict the probability that they are anticancer peptides. As shown in Fig.2a, in terms of AMP generation performance, PepGenWOA comprehensively outperforms all baseline models, achieving lower alignment score and instability score while achieving higher antimicrobial probability. For the best baseline model [36], the alignment score is improved by PepGenWOA by 12.1%, the instability score is improved by 12.6%, and the antimicrobial probability is improved by 5.8%. As shown in Fig.2b, in terms of ACP generation performance, PepGenWOA surpasses all baseline models across the board, including alignment score, instability score and anticancer probability. PepGenWOA exceeds the best baseline model [36] by 14.7% in alignment score, 11.7% in instability score and 8.3% in anticancer probability. The improvements in alignment score indicate that the AMP and ACP sequences generated by PepGenWOA have higher novelty, which means that PepGenWOA illuminates a larger sequence space of therapeutic peptides. The improvements in instability score indicates that the generated sequences are more stable in conformation, which means that PepGenWOA captures the conformational stability that is crucial for AMP and ACP to exert bioactivity. The improvements in antimicrobial or anticancer probability demonstrate that the sequences generated by PepGenWOA have higher probabilities of being therapeutic, highlighting that PepGenWOA understands the bioactive mechanism of antimicrobial or anticancer peptide to a certain extent.

**Fig. 2.**
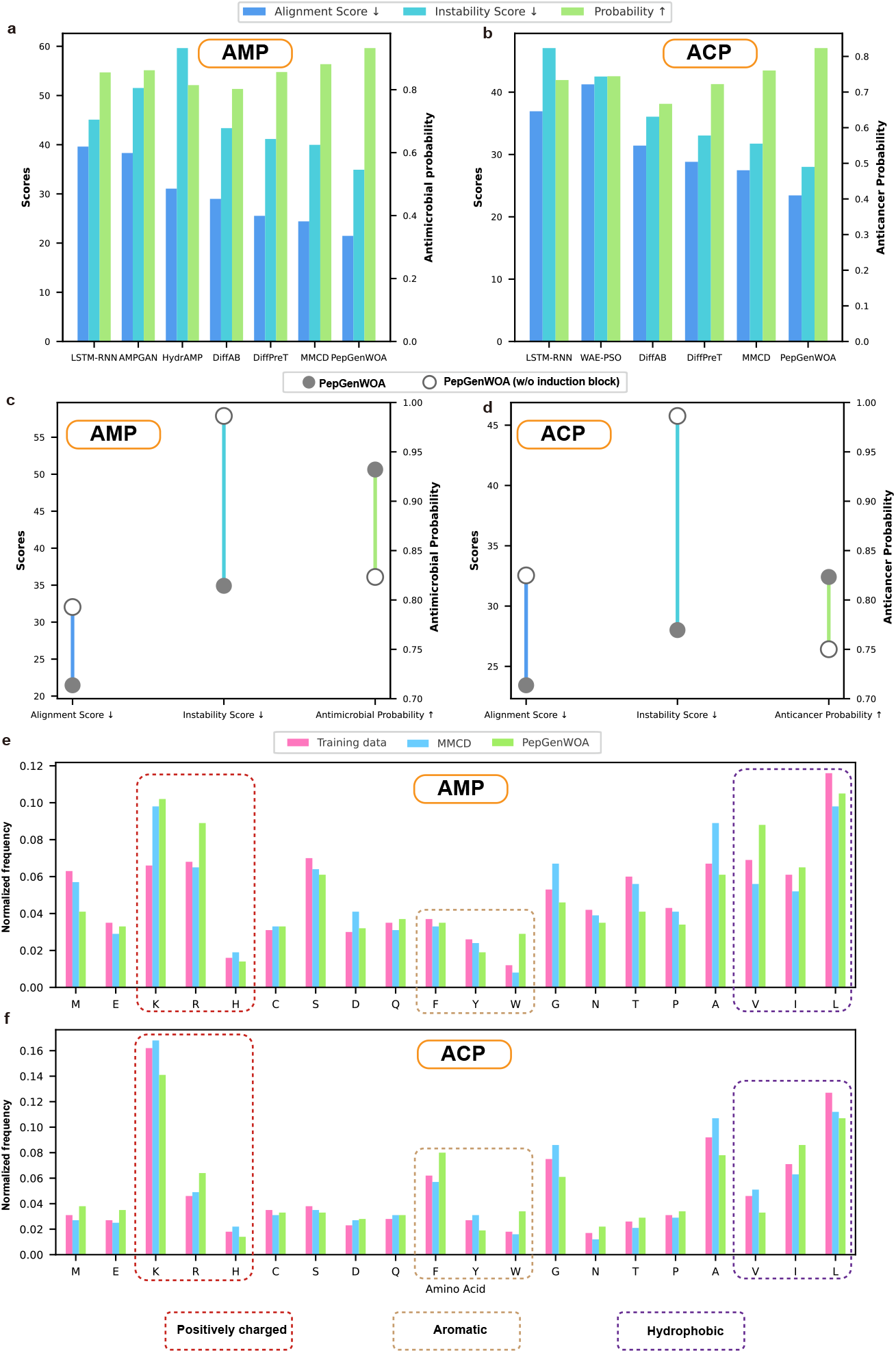
Generation performance for AMP and ACP. **abcd**, Generation performance comparison between different models (ab) and ablation experiments for the proposed induction block (cd). The quality of generated antimicrobial peptides (AMPs, ac) and anticancer peptides (ACPs, bd) is evaluated by three metrics, including alignment score (the left vertical axis), instability score (the left vertical axis), and antimicrobial or anticancer probability (the right vertical axis). The reported results are the average of 1,000 sequences generated by each model. **ef**, Amino acid composition of the generated AMP sequences (e) and ACP sequences (f), where representative positively charged amino acids, aromatic amino acids, and hydrophobic amino acids are highlighted by dashed boxes. The training data, the sequences generated by the best baseline model, and the sequences generated by PepGenWOA are taken separately to calculate the normalized amino acid frequencies.

Second, considering the proposed induction block is a key component of PepGen-WOA, we perform module ablation experiments on AMP and ACP generation tasks to explore its necessity. As shown in Fig.2cd, once the induction block is removed, the generation performance of AMP and ACP is greatly reduced. Specifically, for AMP generation, removing induction block will cause alignment score, instability score, and antimicrobial probability to decay by 49.3%, 65.8%, and 11.7%, respectively. For ACP generation, the decay ratios of the above three metrics are 38.9%, 63.4%, and 8.8%, respectively. Overall, the drastic performance degradation in all three metrics demonstrates that the effectiveness of induction block.

Third, we analyze the generation propensity of the models to explore whether Pep-GenWOA establishes interaction dependencies between residues to capture the shared basic mechanisms of peptide bioactivity. As shown in Fig.2ef, the amino acid compositions of the training data, the generated AMP sequences, and the generated ACP sequences are calculated and normalized. In addition to PepGenWOA, we select the best baseline model MMCD [36] for comparison. From the frequencies of the overall amino acid composition, the compositions of the generated sequences of MMCD and PepGenWOA are roughly consistent with the training data, which shows that these models have learned the probability distribution of amino acid types similar to known AMPs and ACPs. In addition, positively charged amino acids, aromatic amino acids, and hydrophobic amino acids have been shown to be crucial for the activity of AMPs and ACPs by previous studies [58–65]. Specifically, positively charged amino acids can increase the tendency of AMPs or ACPs to interact with negatively charged membranes, enhancing membrane disruption and pore formation [60–63]. Aromatic amino acids can promote the interaction of AMP or ACP with membranes through aromatic ring structures [58, 59]. Hydrophobic amino acids can enhance the ability to bind to and insert into membranes by interacting with the lipid bilayer on the membrane [64, 65]. Therefore, we focus on analyzing the changes in the generation tendency of the above three types of amino acids in different models. We can find that the best baseline model MMCD maintains the amino acid composition close to the training data for the above three types of amino acids, except that the positively charged amino acid K achieves a significant increase in the generation tendency when generating AMPs. In contrast, PepGenWOA achieves a significant increase in the generation tendency for all three types of amino acids, whether generating AMPs or ACPs. For AMP generation, PepGenWOA shows significant growth in the generation tendency for positively charged amino acids K and R, aromatic amino acids W, and hydrophobic amino acids V (Fig.2e). For ACP generation, PepGenWOA shows significant growth in the generation tendency for positively charged amino acids R, aromatic amino acids W and F, and hydrophobic amino acids I (Fig.2f). In summary, PepGenWOA tends to incorporate amino acid types that are beneficial to antimicrobial or anticancer bioactivity during generation, demonstrating that it internalizes the shared basic bioactivity mechanisms of known AMPs and ACPs through unsupervised learning.

### 2.3 Characteristic-guided generation of peptide binders with lifelong learning

Peptide binders have been extensively studied and have become a therapeutic modality with great potential [12, 66–68]. Due to the enormous potential sequence space of peptide binders, wet-lab experimental methods, such as molecular biology-based selection techniques [69, 70] and affinity selection-mass spectrometry [71, 72], have difficulty anchoring a subspace with desired properties. Here, we demonstrate a lifelong learning strategy [73, 74] for incorporating desired characteristics in PepGenWOA by reasonable fine-tuning techniques to achieve characteristic-guided generation of peptide binders (Fig.3a). In this pipeline, we first pretrain PepGenWOA on about 140 million natural short protein sequences. Specifically, considering that peptide binders are generally short in length, these sequences with lengths less than 128 are selected from multiple protein sequence database to promote PepGenWOA to capture patterns of natural short protein sequences, i.e., the world knowledge for peptide generation task (Methods 4.1.2). PepGenWOA pretrained in this way is essentially a peptide-like sequence generator, which is in an unconditional generation state. Subsequently, we hint the desired characteristics, including physicochemical properties and target binding modes, to pretrained PepGenWOA through the fine-tuning strategy. In this section, generating peptide binders with stable conformations that bind SARS-CoV-2 Omicron BA.5 receptor-binding domain (RBD) is adopted as a case study. Two sequence datasets with different characteristics, including a miniprotein scaffold dataset with stable structure [75] and *in silico* synthesized peptides targeting Omicron BA.5 RBD through Rosetta [76], are elaborated for the step-by-step fine-tuning (Methods 4.1.2). Finally, PepGenWOA is expected to perform characteristic-guided conditional generation of peptide binders.

**Fig. 3.**
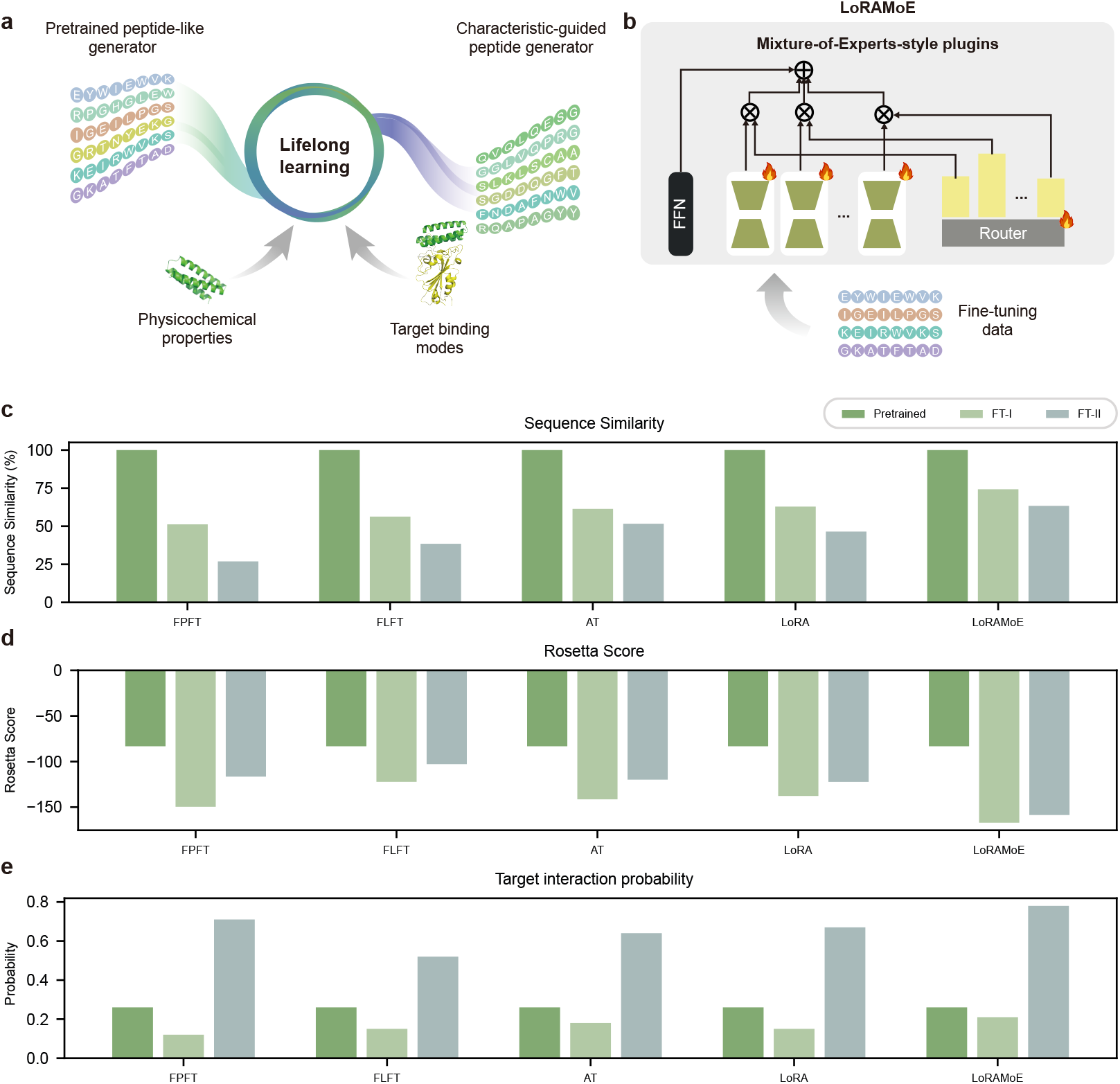
Different fine-tuning strategies for peptide binder generation. **a**, Illustration of characteristic-guided peptide binder generation with lifelong learning. Sequences containing physicochemical properties and target binding modes are adopted to continuously fine-tune the pretrained peptide-like generator to obtain a characteristic-guided peptide generator. **b**, Architecture of LoRAMoE, where multiple low-rank adapters (LoRAs) as adaptable experts (i.e., Mixture-of-Experts-style plugins) are gated into the feed-forward network (FFN) layer of every transformer block by a router. In each finetuning process, only the parameters of the experts and the router are optimized (marked by the flame symbol). **cde**, Evaluation of memory stability and learning plasticity of different fine-tuning strategies, including full-parameter fine-tuning (FPFT), freezing-layer fine-tuning (FLFT), adapter tuning (AT), LoRA, and LoRAMoE. Three metrics, including sequence similarity (c), Rosetta score (d), and target interaction probability (e), are adopted to evaluate the memory stability for world knowledge of natural short protein sequences and the learning plasticity for the new knowledge of structural stability and target binding mode. “Pretrained” denotes the pretrained PepGenWOA, “FT-I” denotes the PepGenWOA fine-tuned by structural stability data, and “FT-II” denotes the PepGenWOA further fine-tuned by target binding mode data. The reported results are the average of 1,000 sequences generated by each model.

For language models, as the scale of fine-tuning data increases, the frozen snapshot of the world learned during pretraining, known as parameterized knowledge [77] or world knowledge [78], will inevitably be destroyed [79]. The above phenomenon is called knowledge forgetting of lifelong learning [80], which leads to the stability-plasticity dilemma [73, 74, 81] between improving the performance of downstream tasks and maintaining world knowledge. In this section, the pretrained PepGenWOA is fine-tuned for multiple rounds with data of different characteristics to introduce new knowledge, facing the severe challenge of knowledge forgetting. To choose a reasonable fine-tuning strategy, five different approaches are implemented to explore the best stability-plasticity trade-off between the basic pattern of natural short protein sequences (i.e., world knowledge) and the desired characteristics as conditional generation constraints (i.e., new knowledge). Specifically, the full-parameter finetuning (FPFT), the fine-tuning approach that freezes the parameters of specific layers (FLFT), the adapter tuning (AT), the low-rank adaptation (LoRA), and the finetuning approach introducing multiple plugins as experts (LoRAMoE) are adopted (Fig.S1 and Fig.3b). Compared with full-parameter fine-tuning, freezing-layer finetuning [82] aims to prevent the destruction of world knowledge by freezing the region storing it. Adapter tuning [83] designs the adapter module added to transformer blocks, which fixes the pretrained model parameters during fine-tuning and only trains the task-specific inserted parameters. LoRA [84] simulates parameter changes through low-rank decomposition, which significantly reduces the number of trainable parameters for downstream tasks. LoRAMoE [85] combines the advantages of LoRA and Mixture of Experts (MoE) [86], where multiple plugins as experts and a router used to gate these experts to assign weights are added to the transformer layers (Methods 4.3).

To determine the best stability-plasticity trade-off, three metrics are adopted, including the sequence similarity, the Rosetta score, and target interaction probability. First, to evaluate the memory stability of different fine-tuning strategies for world knowledge, we adopt the sequence similarity between the sequences generated by the models before and after fine-tuning as the metric for world knowledge forgetting degree. Second, the Rosetta score [76] reflecting the stability of the overall structure is used to evaluate the learning plasticity of the fine-tuned PepGenWOA for the new knowledge of structural stability. Third, a protein-protein interaction prediction model [87] that performs zero-shot predictions is employed to predict the target interaction probability between the generated peptide binders and Omicron BA.5 RBD to evaluate the learning plasticity of the fine-tuned PepGenWOA for the new knowledge of target binding mode.

In this section, the miniprotein scaffold dataset [75] is adopted for the first round of fine-tuning to introduce structural stability, and *in silico* synthesized peptides targeting Omicron BA.5 RBD through Rosetta are adopted for the second round to introduce target binding mode. As shown in Fig.3cde, FPFT leads to the highest world knowledge forgetting degree (i.e., the lowest sequence similarity), but the effective learning of new knowledge led to the suboptimal target interaction probability. FLFT achieves the second highest world knowledge forgetting degree, accompanied by the worst structural stability and target interaction probability, as the difficulty in precisely selecting the frozen parameter region leads to unpredictable fine-tuning benefits. Adapter tuning (AT) and LoRA achieve similar world knowledge forgetting degree, structural stability, and target interaction probability, showing a certain superiority in fine-tuning benefits compared with FPFT and FLFT. LoRAMoE achieves the best results in all three metrics, with 22.8% higher than the suboptimal AT in world knowledge forgetting degree (sequence similarity), 29.7% higher than the suboptimal LoRA in structural stability (Rosetta score), and 6% higher than the suboptimal FPFT in target interaction probability. In addition, we also conduct the experiments under the reversed fine-tuning order, that is, the target binding mode is introduced in the first round and the structural stability is introduced in the second round (Supplementary information Table.S10, Table.S11, Table.S12). The results suggest that LoRAMoE promotes the new knowledge for desired characteristics by preserving the world knowledge for natural short sequence patterns to the greatest extent. Considering that targeting probability is crucial for peptide binders, we finally choose the fine-tuning order of introducing structural stability in the first round and introducing the target binding mode in the second round to maximize the potential for interaction between the generated sequences and the target protein.

### 2.4 Mega-scale structure-based virtual screening of peptide binders

In this section, characteristic-guided PepGenWOA fine-tuned with LoRAMoE initially generates more than 2.2 million peptide binder candidates. It is crucial to efficiently anchor the most promising candidates in the mega-scale synthetic sequence space to reduce the cost of *in vitro* experimental validation. Considering that PepGenWOA is a sequence generation model, we screen candidates from a structural perspective to exclude bias from sequence patterns. Therefore, we introduce a structure-based progressive virtual screening pipeline, including the static structure screening (Fig.4ab) and the binding dynamics screening (Fig.4cd). Considering that directly predicting the complex structures of millions of peptide binders with Omicron BA.5 RBD will result in huge computational overhead, we progressively screen the candidates for their secondary structures, monomer structures, and complex structures with target protein to reduce computing resource requirements and speed up screening processes. First, the secondary structures of the initial generated mega-scale candidates are predicted by ProtTrans [88] to perform a rapid screening with the criterion of 3-helix structure, which is beneficial for structural stability and binding complementarity for Omicron BA.5 RBD [75, 89]. As shown in Fig.4a, the above screening results in the synthetic sequence space being effectively reduced from about 2.2 million to 44,278, which significantly reduces the computational over-head of subsequent monomer and complex structure predictions. Second, the monomer structures are predicted by computationally efficient ColabFold [90] to further reduce the above synthetic space to 6,025. In this screening stage, we calculate the angles and distances between helices to exclude candidates that are difficult to form 3-helix structure. Third, the complex structures between the remaining candidates and Omicron BA.5 RBD are predicted by AF2Complex [91] to assess the binding situations. The analyses for interface residue, targeting atom, and interaction strength are performed to reduce the synthetic sequence space to 157, which is 4 orders of magnitude lower than the initial generation scale. The average structural characteristics of the remained 157 candidates are shown in Table.S13. More importantly, to demonstrate whether static structure-based virtual screening directionally shrinks the synthetic sequence space, we visualize the synthetic space of the remained candidates in different screening stages (Fig.4b). Specifically, we first employ four sequence-based protein descriptors (Methods 4.5.2) to extract comprehensive implicit information of the generated peptide binders. Factor analysis [92] is then adopted for dimension reduction visualization to unveil the changing trend of the synthetic sequence space. The regular contraction of the distribution space corresponding to the three screening stages indicates that static structure screening approximates a subspace of peptide binders with specific characteristics.

Furthermore, binding dynamics screening is performed to identify the most promising candidates based on their binding energies to the target protein. Specifically, the complex structures of the remaining 157 peptide binders with Omicron BA.5 RBD after static structure screening are subjected to molecular dynamics (MD) simulations to estimate their binding energies. To confirm the binding specificity of the generated peptide binders to Omicron BA.5 RBD, we additionally perform MD simulations for the complexes of these 157 candidates with wild-type RBD as negative control. The MM-PBSA method [93] is adopted to calculated the binding energies through two different models for mutual verification, i.e. the non-polar solvation models of solvent-accessible surface area (SASA) and solvent-accessible volume (SAV). The best four binding energies, corresponding to candidates SN01, SN02, SN03, and SN04, are shown in Fig.4c, which are consistent in SASA model and SAV model. With the results of complexes of these candidates with wild-type RBD as negative control, the candidates clearly prefer to target Omicron BA.5 RBD rather than wild-type RBD, demonstrating the binding specificity introduced by target binding mode information in the second round of fine-tuning. In addition, considering the importance of water solubility for the bioactivity of peptide binders, we adopt the Gibbs Free Energy of solvation (*G solvation*) to compare the solubility of the four most promising candidates with two types of RBD. As shown in Fig.4d, each candidate owns a significantly smaller solvent-accessible surface area than the two types of RBD, while *G solvation* is close to or more negative. This results in their specific *G solvation*, i.e. the Gibbs energies of solvation per solvent accessible surface area, being more negative, which indicates that these candidates may have a certain degree of water solubility that is beneficial for peptide biactivity. Finally, we retain the 7 peptide binder candidates with the most negative binding energies for subsequent *in vitro* experimental validation, which is 5 orders of magnitude lower than the initial generation scale.

**Fig. 4.**
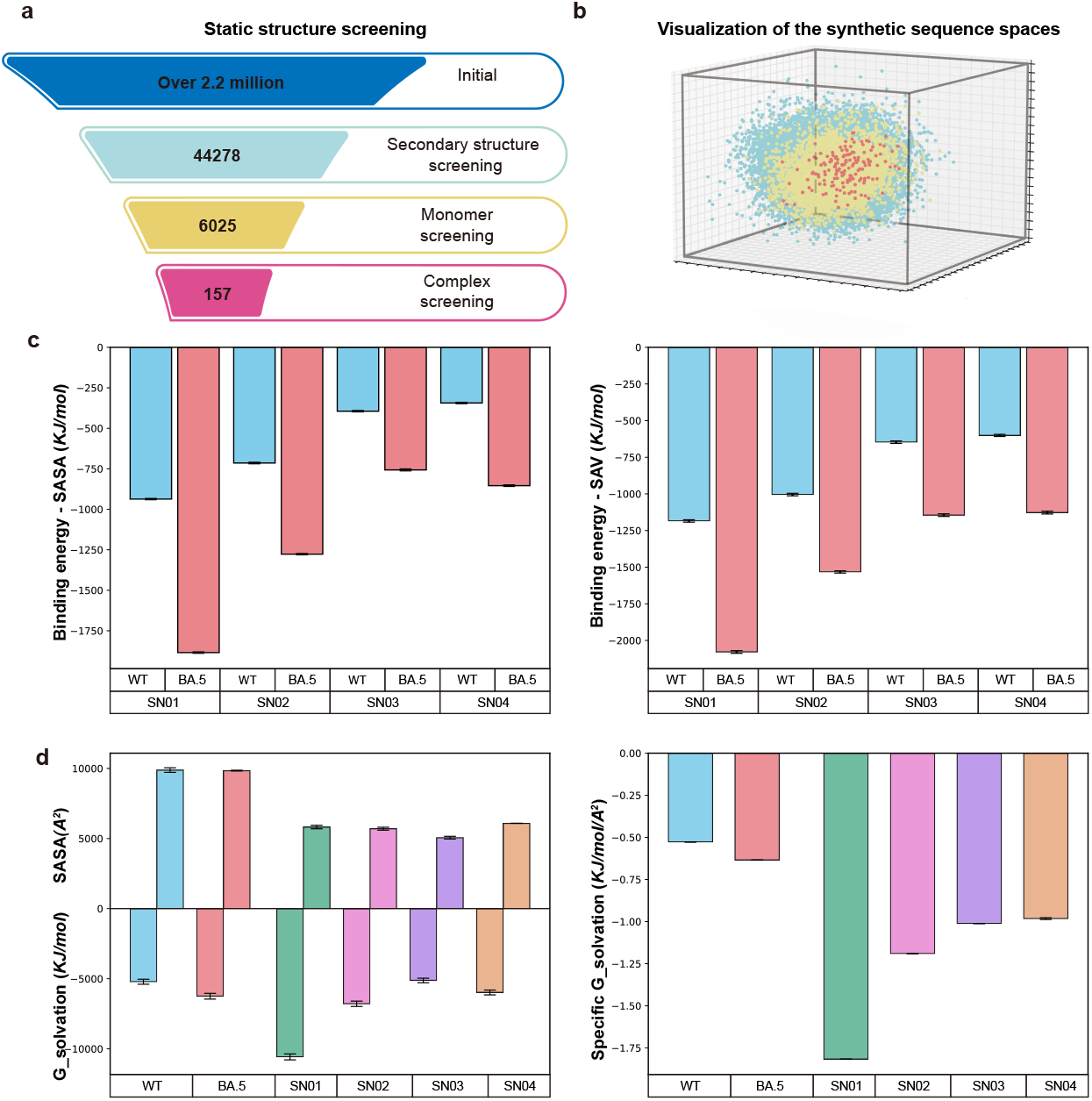
Mega-scale structure-based virtual screening. **a**, The illustration of static structure virtual screening. The generated peptide binders are screened by diverse criteria of three levels of static structure, including secondary structure, monomer, and complex. The number of candidates is annotated corresponding to the different screening stages for static structures. “Initial” denotes the initially stage, “secondary structure screening”, “monomer screening”, and “complex screening” denote progressive screening stages. **b**, The visualization of the synthetic sequence spaces at different screening stages for static structures. The data points are colored according to the different screening stages. **c**, Binding energies of the four most promising candidates to the target estimated by molecular dynamics simulations, in which the non-polar solvation models of solvent-accessible surface area (SASA, left panel) and solvent-accessible volume (SAV, right panel) are adopted for mutual verification. “BA.5” denotes that the target protein is Omicron BA.5 RBD, and “WT” denotes that the target protein is wild-type RBD, which is a negative control to demonstrate binding specificity. **d**, Solvation free energy (G solvation, left panel) and corresponding specific G solvation (the Gibbs energies of solvation per solvent accessible surface area, right panel) of the four most promising candidates. Omicron BA.5 RBD and wild-type RBD are adopted to compare their water solubility with the candidates.

### 2.5 *In vitro* experimental validation of screened peptide binders

To verify the target binding ability of the screened peptide binders, yeast surface display and flow cytometry experiments are preformed, in which the nanobody Ty1 [94] targeting wild-type RBD is used as a positive control (Methods 4.6). The apparent binding affinities of the candidates targeting Omicron BA.5 RBD or wild-type RBD are determined by flow cytometry, where candidates with binding affinity at the *µ*M level are considered positive samples with successful binding (Supplementary information Table.S19). As shown in Fig.5a, PepGwnWOA achieves a binding rate of 28.6% targeting Omicron BA.5 RBD, while the binding rate targeting wild-type RBD is 0% (Supplementary information Table.S20), indicating that the target binding mode introduced by the fine-tuning strategy of Mixture-of-Experts-style plugins improves the binding specificity of the generated peptide binders. The display efficiency and target binding of the positive control Ty1 and the two candidates that successfully bind to Omicron BA.5 RBD are imaged by confocal microscopy in Fig.5bc. Overall, the above *in vitro* experimental validation results demonstrate that characteristic-guided PepGenWOA and mega-scale structure-based virtual screening constitute an low-cost generative design pipeline for peptide binders, which efficiently anchors the bioactive subspace in the mega-scale sequence space.

**Fig. 5.**
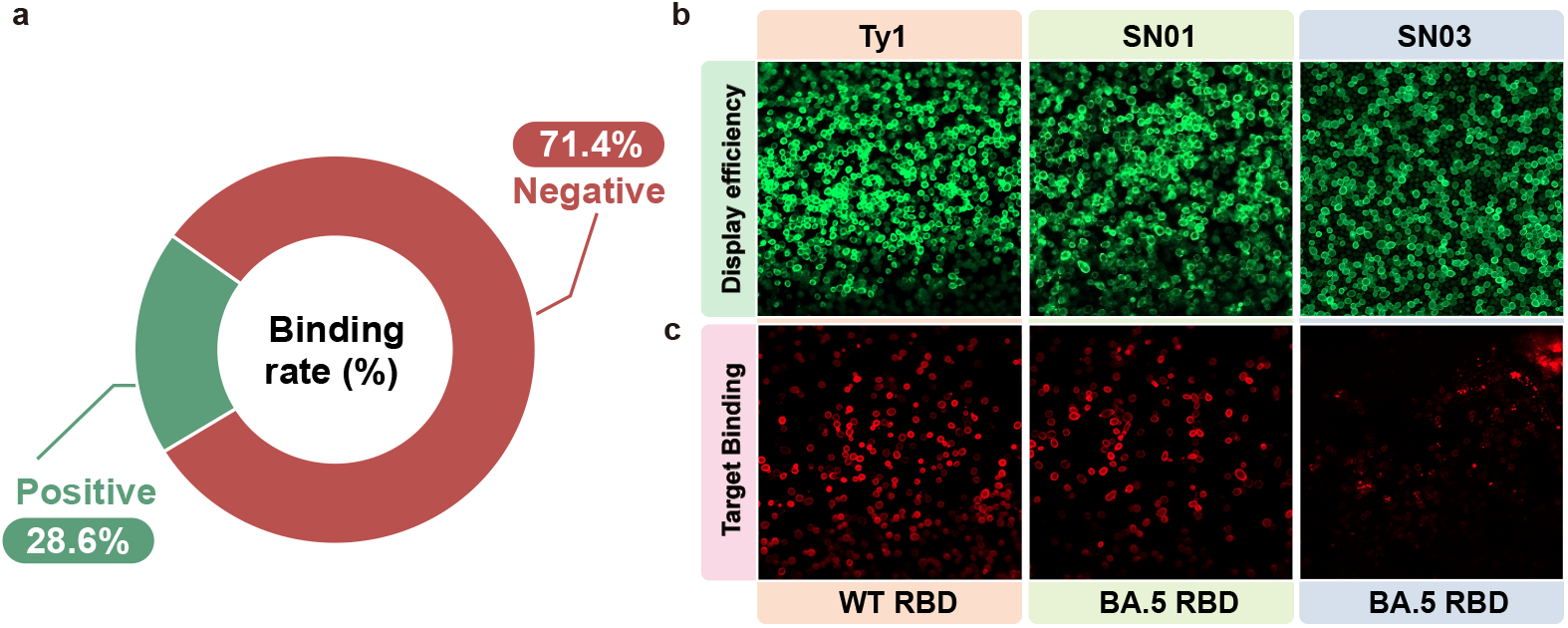
Experimental validation of screened peptide binders. The nanobody Ty1 that binds to wild-type RBD is adopted as a positive control. **a**, The 28.6% binding rate of the candidates targeting Omicron BA.5 RBD, which is evaluated by yeast surface display and flow cytometry. **bc**, The display efficiency (a) and target binding (b) of the positive control Ty1 (left panel) and the two candidates (SN01, middle panel; SN03, right panel) that successfully bind to Omicron BA.5 RBD imaged by confocal microscopy. “WT RBD” denotes the wild-type RBD, and “BA.5 RBD” denotes the Omicron BA.5 RBD.

## 3 Discussion

By introducing tolerance for out-of-order inputs as an inductive bias of the autoregressive architecture, PepGenWOA not only predict the next residue during training, but also understand and build the accurate interaction dependencies between residues that determine the peptide bioactivity. The generation performance of AMP and ACP comprehensively outperforms the competitors, demonstrating a significant propensity to incorporate specific types of residues that are beneficial for antimicrobial or anticancer bioactivity. With the lifelong learning paradigm, PepGenWOA is first pretrained to capture the world knowledge of natural short protein sequence patterns, and then fine-tuned by Mixture-of-Experts-style plugins for the best trade-off between memory stability for world knowledge and learning plasticity for new downstream tasks. The well fine-tuned PepGenWOA initially generated more than 2.2 million peptide binders, which are reduced by 5 orders of magnitude by mega-scale structure-based virtual screening to efficiently anchor 7 candidates for *in vitro* experimental validation. A target binding rate of 28.6% with binding specificity further confirms the effectiveness and low cost of the characteristic-guided generation pipeline of peptide binders.

Although PepGenWOA is a promising unified framework for generative peptide design, it still has limitations in terms of controllable generation, which is caused by data scarcity. The number of natural peptide sequences with specific physicochemical properties or target binding modes is often very limited, which makes it difficult to effectively fine-tune the language model, even if multiple parameter efficient fine-tuning strategies are adopted. The synthetic data strategy based on Rosetta software is adopted in this work, but the synthesis quality is still unknown, so further development of data enhancement methods for generative peptide design is necessary. In the future, as the scale of natural or high-quality synthetic peptide data gradually increases, the controllable generation ability of PepGenWOA is expected to be further improved, thereby achieving *de novo* design with higher targeting binding success rate.

## 4 Methods

### 4.1 Datasets

#### 4.1.1 AMP and ACP dataset

For the AMP and ACP generation tasks, PepGenWOA is trained with compiled sequence datasets, including 20,129 AMP sequences and 4,381 ACP sequences, of a previous study [36]. Specifically, the AMP dataset is integrated from 7 open-source AMP databases, including LAMP [95], YADAMP [96], SATPdb [97], DRAMP [98], DBAMP [99], CAMP [58], and APD3 [100]. Similarly, the ACP dataset is also integrated from 6 open-source AMP databases, including CancerPPD [101], LAMP [95], DRAMP [98], DBAMP [99], SATPdb [97], and APD3 [100].

#### 4.1.2 Peptide binder dataset

##### Pretraining dataset

In order to capture the patterns of natural short protein sequences, we screen natural protein sequences with lengths less than 128 from three protein sequence databases, including BFD^1^, Pfam^2^, and UniProt^3^, for the pretraining of PepGenWOA. Finally, about 140 million raw protein sequences are retained.

##### Fine-tuning dataset

For the fine-tuning introducing structural stability, a miniprotein scaffold dataset with stable structure [75] is adopted. For the fine-tuning introducing target binding mode, *in silico* synthesized peptides targeting Omicron BA.5 RBD through Rosetta [76] are adopted. Specifically, we employ Rosetta to constrain the angiotensin-converting enzyme 2 (ACE2) interface in contact with the Omicron BA.5 RBD to generate a variety of potentially targeting peptides. First, the complex structure of Omicron BA.5 RBD with ACE2 (PDB ID: 7XWA) [102] is refined with the Rosetta FastRelax protocol. Second, the interface of ACE2 (residue 23-46) interacting with RBD is extracted for subsequent peptide generation. Third, the ACE2 helix (residue 23-46) is adopted as one of the helices to design a variety of 3-helical bundles by the Rosetta Remodel blueprint [103], where thousands of blueprints are generated with different lengths of the helices and the loop types. Fourth, backbones are generated by Rosetta Monte Carlo-based fragment assembly protocol [104], where Omicron BA.5 RBD structure is loaded into a hashing grid to ensure the generated backbones target it. Fifth, interface residues on the backbones except for the ACE2 helix part are designed to make more contact with RBD by the Rosetta PackRotamersMover. Sixth, the whole peptide besides the main interaction residues on the ACE2 helix is designed by the Rosetta FastDesign protocol with side-chain rotamer optimization and energy minimization. Seventh, the whole peptide is optimized with more Rosetta FastDesign protocols to improve shape and chemical complementarity. Through the above process, we finally synthesize more than 160,000 peptide sequences with the potential to target Omicron BA.5 RBD.

### 4.2 Pretraining configuration for peptide binder generation

The detailed pretraining configuration for peptide binder generation is described as follows. The number of layers is 24 (including the induction block), the hidden layer size is 1,024, the intermediate size of the multilayer perceptron (MLP) block is 4,096, and the number of attention heads is 16. Therefore, the total number of model parameters is approximately 350 million. The adopted optimizer is Adam, the weight decay is 0.01, and the learning rate is 1e-4. The Cross-Entropy loss function is adopted and cosine decay is used for the learning rate scheduling. PepGenWOA is pretrained across 128 Neural Processing Units (NPUs) for about ten hours.

### 4.3 Fine-tuning with LoRAMoE for peptide binder generation

LoRAMoE [85] is Mixture-of-Experts-style plugin, which combines the advantages of LoRA [84] and Mixture of Experts (MoE) [86]. A router network is adopted to integrate several experts (i.e., LoRAs), thereby enhance the model’s learning plasticity for downstream tasks while alleviating the world knowledge forgetting.

The forward propagation process of feed-forward neural (FFN) network and the matrix operation of the linear layer in the traditional transformer architecture are as follows:

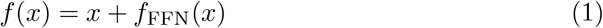

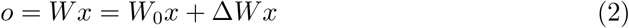

where 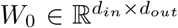 denotes the parameter matrix of the backbone model and 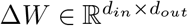 denotes the updated parameter during the training process.

LoRAMoE replaces the linear layer in the FFN with the MoE-style LoRA, where the backbone model is frozen during training and only Δ*W* is updated. After replacing the parameter matrix of the experts with LoRA, the forward process of the LoRAMoE layer can be written as equation.5:

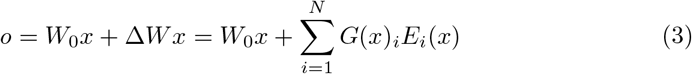

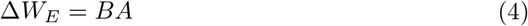

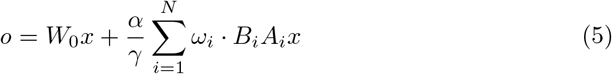

In equation.3, *E*_*i*_(*·*) denotes the *i*-th expert, *G*(*·*) = Softmax(xW_g_) denotes the router, *W*_*g*_ denotes the trainable parameter matrix of the router network. In equation.4, the rank *γ ≪ min*(*d*_*in*_, *d*_*out*_), 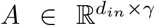, and 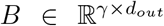. In equation.5, *ω*_*i*_ and *α* represent the *i*-th expert weight and the constant hyperparameter, respectively.

### 4.4 Evaluation metrics

#### 4.4.1 AMP and ACP evaluation

Alignment score is calculated with the PairwiseAligner of Biopython tool [105], where the BLOSUM62 is adopted as the alignment scoring matrix. Instability score is calculated with PeptideDescriptor of the modlAMP tool [57]. The antimicrobial probability of AMP is predicted by the CAMP model [58], and the anticancer probability of ACP is predicted by the AntiCP model [59].

#### 4.4.2 Peptide binder evaluation

The sequence similarity adopts the normalized alignment score between 0 and 1. The Rosetta score is obtained with the score function of Rosetta software. The target interaction probability is predicted by a protein-protein interaction prediction model [87].

### 4.5 Structure-based virtual screening of peptide binders

#### 4.5.1 Static structure screening

First, for the secondary structure screening, the criterion of 3-helix structure is adopted referring to previous studies [75, 89]. Second, for the monomer structure screening, the angle and distance between helices are calculated to exclude candidates that are difficult to form 3-helix structure. Specifically, considering that too large a helix angle or helix distance results in failure to form 3-helix structure, we set screening thresholds for the helix angle and distance. The helix angle reflecting the parallelism between the helices is required to be less than 20, and the helix distance representing the average centroid distance of helices is required to be less than 12Å. Third, for the complex structure screening, the interface residue, targeting atom, and interaction strength are analyzed to comprehensively evaluate the binding between the peptide binder and the target protein. Specifically, the interface residue representing the number of residues at the binding interface of the complex structure is required to greater than 60. Targeting atom represents the number of backbone atoms in the peptide binder that are spatially close to the target, which is required to greater than 40. In addition, the number of miss-targeting atoms is required to less than 200. Here, the number of hydrogen bonds in the complex structure is adopted to evaluate the interaction strength, which is required to greater than 5.

#### 4.5.2 Visualization of the synthetic sequence space

To demonstrate whether static structure-based virtual screening directionally shrinks the synthetic sequence space, we visualize the synthetic space of the remained candidates in different screening stages with sequence-based protein descriptors and factor analysis. Considering that combining multiple protein descriptors allows us to gain a more comprehensive representation of the underlying patterns of peptide binder sequences, we employ the following multiple length-independent descriptors to ensure that sequences of different lengths have equal-length descriptor vectors. All the descriptors are finally concatenated together for factor analysis.

##### k-mer spectra

A *k*-mer refers to a set of short sub-sequences containing several amino acids within a protein sequence, which is defined as:

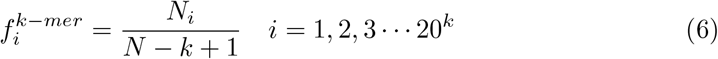

Where *N*_*i*_ denotes the number of the *i*-th sub-sequence of length *k*, and *N* is the length of the current sequence. We employ *k* values of 1, 2, and 3, which result in 20 (20 standard amino acids), 400 (20^2^), and 8000 (20^3^) dimensions, respectively, for the generated *k*-mer spectra.

##### Normalized Moreau-Broto Autocorrelation descriptor

Normalized Moreau-Broto Autocorrelation (NMBAC) descriptors [106] capture the local correlation of physicochemical properties within a protein by measuring the interaction between amino acids. The general formula for Normalized Moreau-Broto Autocorrelation is as follows:

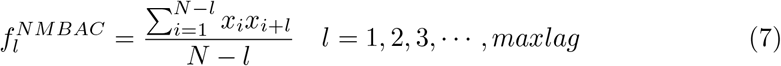

Where *x*_*i*_ and *x*_*i*+*l*_ represent the normalized physicochemical property value of the *i*-th and *i* + *l*-th amino acid in the protein sequence, *maxlag* is the max lag between amino acids being compared, which is set to 30 here. We considered the following eight properties for the NMBAC calculation: Hydrophobicity, Average Flexibility, Polarizability, Free Energy, Residue Accessible Surface Area (ASA), Residue Volume, Steric, and Mutability, which generates an 8 *×* 30 = 240-dimensional vector.

##### Quasi-sequence-order descriptor

Quasi-sequence-order (QSO) descriptors [107] have been widely used in protein sequence analysis, structural prediction, and protein-protein interaction methods. The first 20 quasi-sequence-order descriptors can be calculated by:

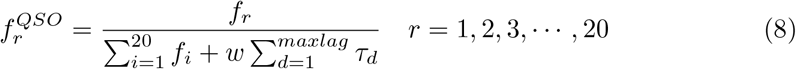

Where *f*_*i*_ is the occurrence of amino acid *i* within the sequence, and *w* is a weight factor. The remaining descriptors are defined as:

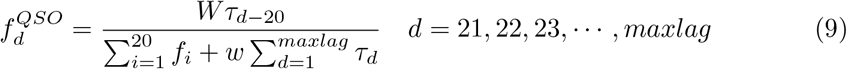

Both Schneider-Wrede physicochemical and Grantham chemical distance matrices are employed here. We set *maxlag* to 30, which generates 100 dimensions.

##### Composition, Transition, and Distribution descriptor

The Composition, Transition, and Distribution (CTD) descriptor [108] is a sequence-based descriptor that represents the amino acid sequence of a protein in terms of its physicochemical properties. The CTD descriptor is composed of three components: Composition (C), Transition (T), and Distribution (D). In this study, we only focus on the Distribution component, as it provides complementary information to the other descriptors used in our study. Specifically, we first divide the amino acids into three classes according to their Polarity. Subsequently, for each class, we calculate five (1%, 25%, 50%, 75%, and 100% residues) distribution descriptors of the whole sequence. Thus, a total of 15 descriptors are obtained.

#### 4.5.3 Binding dynamics screening

GROMACS 2021 [109], OpenMM 7.5 [110] software package, AMBER99SB-ildn force-field [111], and TIP3P water model are adopted for all the molecular dynamics simulations, with a time step of 2 fs. Specifically, first, 1,000 steps of minimization are carried out. Second, for complete relaxation, 4 ns pre-equilibration simulations with restrained coordinates of the heavy atoms are performed. Third, a production simulation with an isothermal-isobaric (NPT) ensemble at 1 atm and 300 K is conducted for a duration of 500 ns. Fourth, the last 10 ns of each trajectory are used for MM-PBSA calculations [93] through the g mmpbsa tools [112] in GROMACS, where 100 snapshots every 100 ps of each trajectory are adopted.

### 4.6 *In vitro* experimental validation

#### 4.6.1 Protein expression and purification

Omicron BA.5 RBD (R319-F541) are expressed with Bac-to-Bac Baculovirus Expression System (Invitrogen). The corresponding genes are synthesized and subcloned into a modified pFastBac1 vector (Invitrogen), which contains an N-terminal GP67 signal peptide sequence and a C-terminal 6xHis purification tag. The expressing construct is transformed into bacterial DH10bac competent cell, the recombinant bacmid is then extracted and transfected into Sf9 insect cell with Cellfectin II reagent (Invitrogen). After two-rounds of amplification, the recombinant baculovirus with high-titer are harvested and mixed with Hi5 insect cells (2 *×* 10^6^ cells per mL). After 60 hours of infection, the cell culture containing the secreted proteins is harvested. Protein is purified with Ni-column (HiTrap Excel, GE Healthcare) and gel filtration column (Superdex 200, GE Healthcare), and the purified protein is concentrated and stored in *−*80^*°*^*C*. PBS buffer is used for all purification steps.

#### 4.6.2 Yeast display vectors construction

pCT2UnaG is a derivative of yeast surface display vector pCTCon2 [113]. A new display cassette is constructed and encodes a leader sequence (LS: appS4), which is followed by a multi cloning site (MCS), a flexible linker sequence, the Aga2p anchor protein, and an UnaG protein. The original cassette in the PCTcon2 vector is replaced by the new one with homologous recombination. All the designed peptides are synthesized and cloned between NdeI and BamHI site for yeast surface display. Yeast is grown in SDCAA medium before induction in SGCAA medium. After induction for 12–18 hours, expression media is supplemented with 5 nM bilirubin. After one hour incubation, cells are collected (3000 g, 5 min) and washed in chilled display buffer PBSA (50 mM NaPO4 pH 8, 20 mM NaCl, 0.5% BSA), and then subjected to analysis.

#### 4.6.3Display and binding determination by flow cytometry

Cells are incubated with varying concentrations of wild type RBD or Omicron BA.5 RBD. After approximately one hour, cells are washed again in a chilled buffer and then incubated on ice. The antibody-based labeling procedure is performed with His-tag Antibody and Alex647 conjugated His antibody. Cells displaying the nanobody protein-Ty1, which binds the wild-type RBD, are used as the positive control and the unlabled yeast cells are used as negative control. For fluorescence activated cell sorting (FACS) analysis of each peptide binder, the previously antibody-labeled cells (10^6^) are transferred to a tube in a final volume of 300 µl with PBSA buffer and applied on a LSRFortessa flow cytometers (BD LSRFortessa, USA).

For each sample, 10,000 yeast cells are analyzed with FlowJo sofware (FlowJo, LLC). The apparent affinities of the positive control and selected peptide binders displayed on yeast are determined by flow cytometry. With a serial dilution, the mean fluorescence intensity (MFI) of each concentration is recorded and plotted against the concentration of the target protein. The one-site specific binding model in Origin2023 is used to calculated the apparent *K*_*D*_.

#### 4.6.4 Confocal microscopy

Yeast cells prepared as above are used for confocal microscopy imaging. Under a confocal laser scanning microscope (Zeiss, LSM980, Germany), cells are observed with excitation wavelength of 543 nm and 647 nm using 50-fold magnifying objective.

Key Points

- We propose a weakly order-dependent autoregressive language modeling architecture, PepGenWOA, for bioactive peptide generative design with tolerating out-of-order input as inductive bias.
- PepGenWOA not only predicts the next residue during unsupervised training on massive protein sequences, but also understands the correct semantics of the random residue order, i.e., the correct interaction dependencies between residues that determine the peptide bioactivity.
- The generation performance of antimicrobial peptide and anticancer peptide comprehensively outperforms other methods, demonstrating a significant propensity to incorporate specific types of residues that are beneficial for antimicrobial or anticancer bioactivity.
- With the lifelong learning paradigm, PepGenWOA is fine-tuned for characteristic-guided peptide binder generation, and subsequent mega-scale virtual screening reduces the number of candidates for *in vitro* experimental validation by 5 orders of magnitude, achieving a target binding rate of 28.6% with binding specificity.
- PepGenWOA represents a unified architecture for peptide generative design that can be flexibly customized for different task requirements, thereby harnessing generative language models to reach the “dark space” of biological language.

## Data availability

The antimicrobial peptide and anticancer peptide dataset is publicly available from a previous study [36]. The pretraining dataset for peptide binder generation can be obtained from BFD (https://bfd.mmseqs.com/), Pfam (http://pfam.xfam.org/) and UniProt (https://www.uniprot.org/). The miniprotein scaffold dataset for fine-tuning is also publicly available from a previous study [75] (https://files.ipd.uw.edu/pub/robust_de_novo_design_minibinders_2021/supplemental_files/scaffolds.tar.gz).

## Code availability

Relevant code and models are available via GitHub at https://github.com/ZhiweiNiepku/PepGenWOA.

## Supplementary information

### S1 Baseline model

PepGenWOA is compared with the baseline models for AMP generation and ACP gen-958 eration. All baseline models are implemented with the original settings in their papers. 959 Specifically, for AMP generation, the LSTM-RNN [22], AMPGAN [29], HydrAMP 960 [28], DiffAB [38], DiffPreT [37], and MMCD models [36] are implemented for comparison. For ACP generation, the LSTM-RNN [22], WAE-PSO [27], DiffAB [38], DiffPreT [37], and MMCD [36] are implemented for comparison.

### S2 Generation performance of AMP and ACP

**Table S1.** Generation performance of AMP.

| Model | Alignment score ↓ | Instability score ↓ | Antimicrobial probability ↑ |
| --- | --- | --- | --- |
| LSTM-RNN | 39.6164 | 45.0862 | 0.8550 |
| AMPGAN | 38.3080 | 51.5236 | 0.8617 |
| HydrAMP | 31.0662 | 59.6340 | 0.8145 |
| DiffAB | 28.9849 | 43.3607 | 0.8024 |
| DiffPreT | 25.5385 | 41.1629 | 0.8560 |
| MMCD | 24.4107 | 39.9649 | 0.8810 |
| PepGenWOA(Ours) | 21.4672 | 34.9121 | 0.9321 |

**Table S2.** Generation performance of ACp.

| Model | Alignment score ↓ | Instability score ↓ | Anticancer probability ↑ |
| --- | --- | --- | --- |
| LSTM-RNN | 36.9302 | 47.0669 | 0.7336 |
| WAE-PSO | 41.2524 | 42.5061 | 0.7443 |
| DiffAB | 31.4220 | 36.0610 | 0.6669 |
| DiffPreT | 28.8245 | 33.0405 | 0.7222 |
| MMCD | 27.4685 | 31.7381 | 0.7604 |
| PepGenWOA(Ours) | 23.4361 | 28.0121 | 0.8233 |

### S3 Ablation experiments for induction block

**Table S3.** Ablation results on AMP.

| Model | Alignment score ↓ | Instability score ↓ | Antimicrobial probability ↑ |
| --- | --- | --- | --- |
| PepGenWOA(w/o induction block) | 32.0455 | 57.8801 | 0.8233 |
| PepGenWOA(Original) | 21.4672 | 34.9121 | 0.9321 |

**Table S4.** Ablation results on ACP.

| Model | Alignment score ↓ | Instability score ↓ | Anticancer probability ↑ |
| --- | --- | --- | --- |
| PepGenWOA(w/o induction block) | 32.5437 | 45.7656 | 0.7501 |
| PepGenWOA(Original) | 23.4361 | 28.0121 | 0.8233 |

### S4 Amino acid composition of generated AMPs and ACPs

**Table S5.**
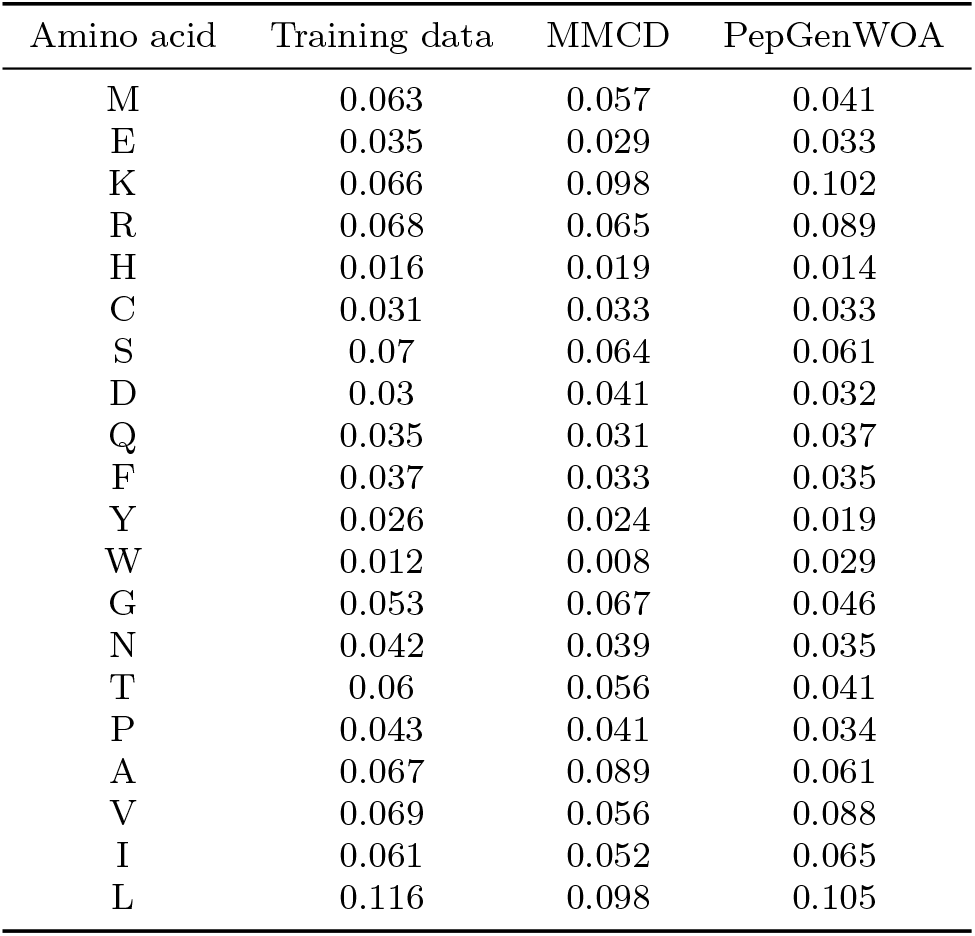
Amino acid composition of generated AMPs.

| Amino acid | Training data | MMCD | PepGenWOA |
| --- | --- | --- | --- |
| M | 0.063 | 0.057 | 0.041 |
| E | 0.035 | 0.029 | 0.033 |
| K | 0.066 | 0.098 | 0.102 |
| R | 0.068 | 0.065 | 0.089 |
| H | 0.016 | 0.019 | 0.014 |
| C | 0.031 | 0.033 | 0.033 |
| S | 0.07 | 0.064 | 0.061 |
| D | 0.03 | 0.041 | 0.032 |
| Q | 0.035 | 0.031 | 0.037 |
| F | 0.037 | 0.033 | 0.035 |
| Y | 0.026 | 0.024 | 0.019 |
| W | 0.012 | 0.008 | 0.029 |
| G | 0.053 | 0.067 | 0.046 |
| N | 0.042 | 0.039 | 0.035 |
| T | 0.06 | 0.056 | 0.041 |
| P | 0.043 | 0.041 | 0.034 |
| A | 0.067 | 0.089 | 0.061 |
| V | 0.069 | 0.056 | 0.088 |
| I | 0.061 | 0.052 | 0.065 |
| L | 0.116 | 0.098 | 0.105 |

**Table S6.**
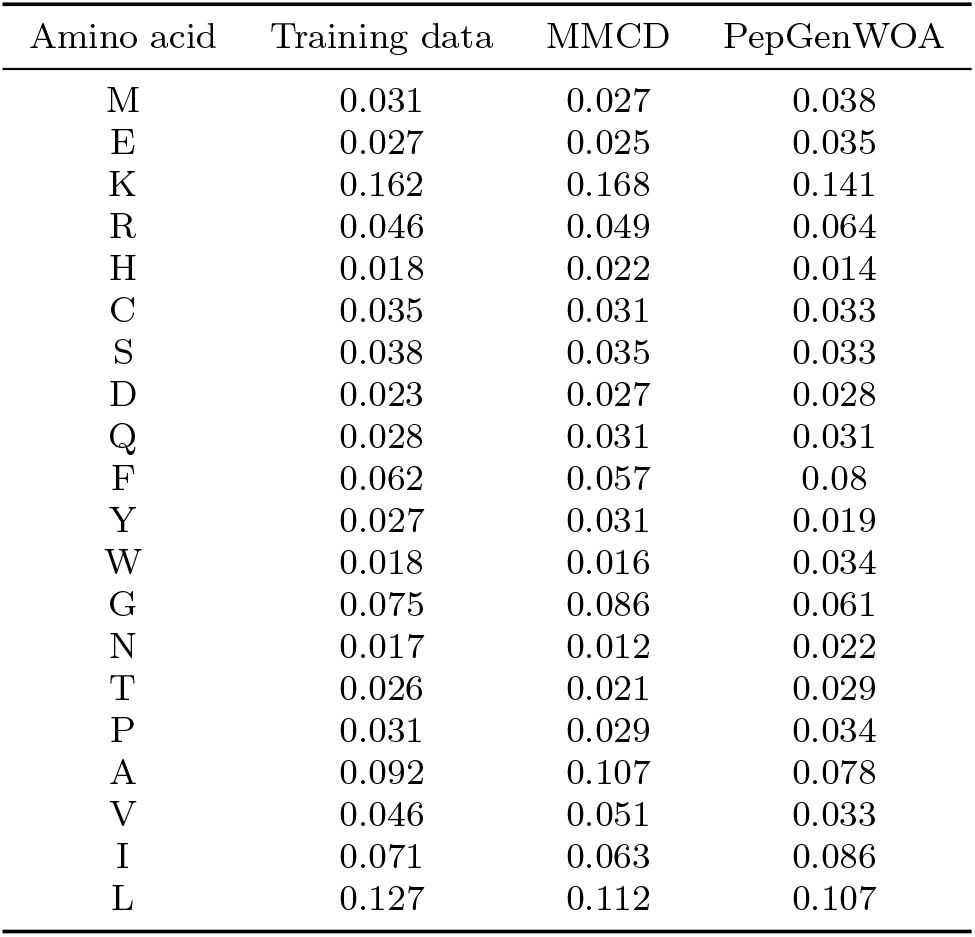
Amino acid composition of generated ACPs.

### S5 Evaluation of different fine-tuning strategies

**Fig. S1.**
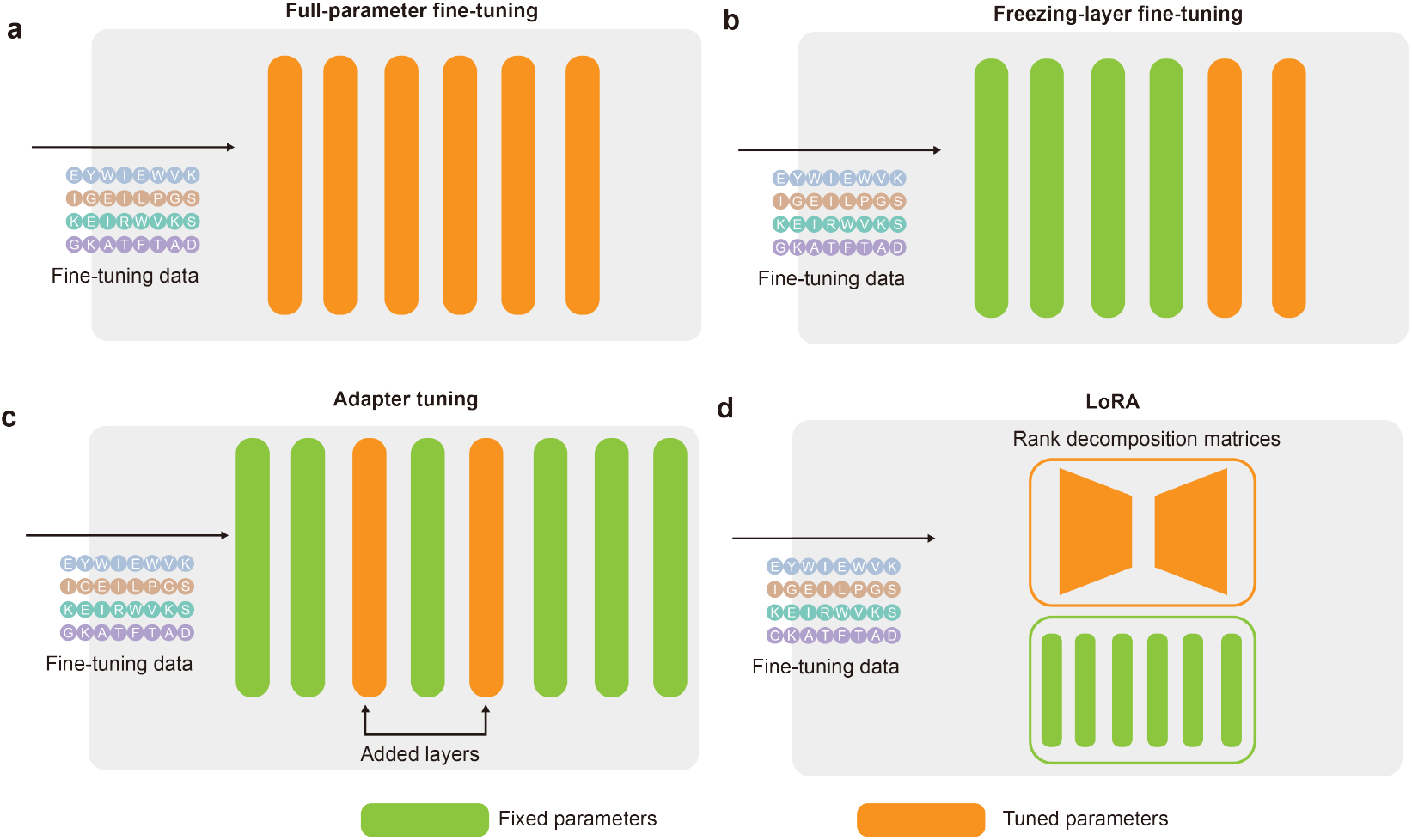
Illustration of different fine-tuning strategies. Full-parameter fine-tuning updates all parameters of the model to adapt to a specific downstream task (a), while freezing-layer fine-tuning selectively updates some parameters, where the region storing world knowledge is frozen (b). Adapter tuning adds the adapter layers to transformer blocks, fixing the pretrained model parameters during fine-tuning for the task-specific inserted parameters updating (c). LoRA adopts the low-rank decomposition to simulate parameter changes, thereby significantly reducing the number of trainable parameters for downstream tasks (d).

**Table S7.** Sequence similarity of different fine-tuning strategies.

| Strategies | Pretrained | FT-I | FT-II |
| --- | --- | --- | --- |
| Full-parameter fine-tuning | 100% | 51.23% | 26.89% |
| Freezing-layer fine-tuning | 100% | 56.33% | 38.46% |
| Adapter tuning | 100% | 61.36% | 51.62% |
| LoRA | 100% | 62.89% | 46.49% |
| LoRAMoE | 100% | 74.26% | 63.38% |

**Table S8.** Rosetta score of different fine-tuning strategies.

| Strategies | Pretrained | FT-I | FT-II |
| --- | --- | --- | --- |
| Full-parameter fine-tuning | -83.24 | -149.63 | -116.53 |
| Freezing-layer fine-tuning | -83.24 | -122.27 | -102.89 |
| Adapter tuning | -83.24 | -141.43 | -119.88 |
| LoRA | -83.24 | -137.78 | -122.33 |
| LoRAMoE | -83.24 | -167.05 | -158.67 |

**Table S9.**
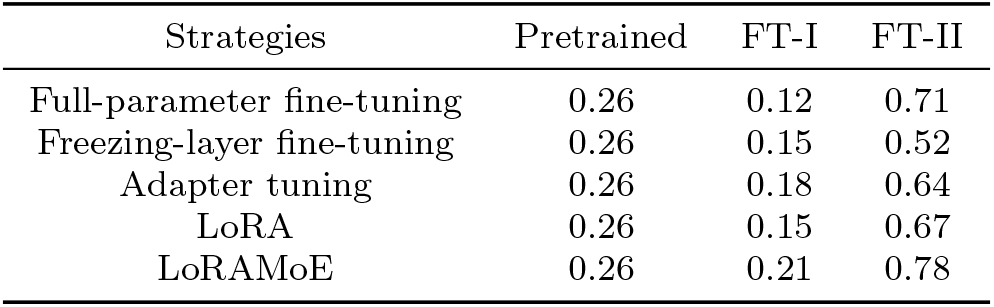
Target interaction probability of different fine-tuning strategies.

| Strategies | Pretrained | FT-I | FT-II |
| --- | --- | --- | --- |
| Full-parameter fine-tuning | 0.26 | 0.12 | 0.71 |
| Freezing-layer fine-tuning | 0.26 | 0.15 | 0.52 |
| Adapter tuning | 0.26 | 0.18 | 0.64 |
| LoRA | 0.26 | 0.15 | 0.67 |
| LoRAMoE | 0.26 | 0.21 | 0.78 |

**Table S10.** Sequence similarity of different fine-tuning strategies under the reversed fine-tuning order.

| Strategies | Pretrained | FT-I | FT-II |
| --- | --- | --- | --- |
| Full-parameter fine-tuning | 100% | 75.54% | 20.17% |
| Freezing-layer fine-tuning | 100% | 79.38% | 37.84% |
| Adapter tuning | 100% | 84.33% | 42.32% |
| LoRA | 100% | 81.72% | 51.89% |
| LoRAMoE | 100% | 90.13% | 60.14% |

**Table S11.**
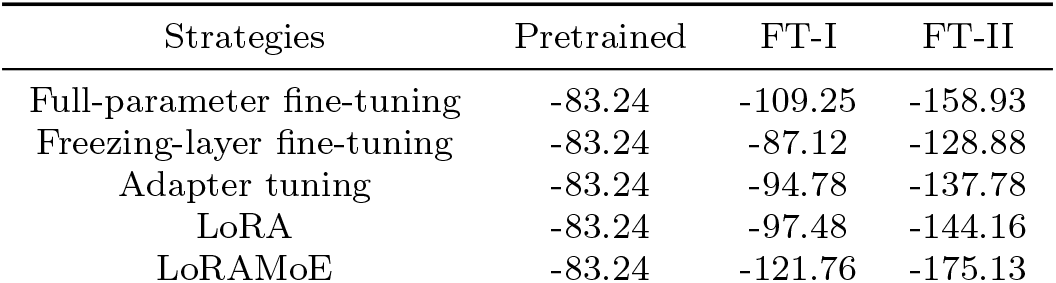
Rosetta score of different fine-tuning strategies under the reversed fine-tuning order.

| Strategies | Pretrained | FT-I | FT-II |
| --- | --- | --- | --- |
| Full-parameter fine-tuning | -83.24 | -109.25 | -158.93 |
| Freezing-layer fine-tuning | -83.24 | -87.12 | -128.88 |
| Adapter tuning | -83.24 | -94.78 | -137.78 |
| LoRA | -83.24 | -97.48 | -144.16 |
| LoRAMoE | -83.24 | -121.76 | -175.13 |

**Table S12.**
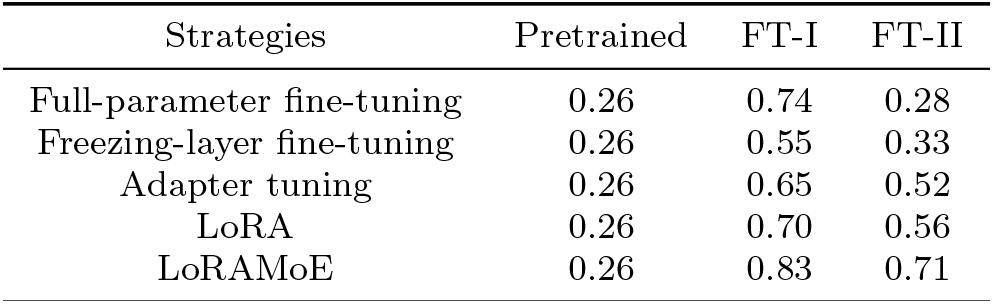
Target interaction probability of different fine-tuning strategies under the reversed fine-tuning order.

| Strategies | Pretrained | FT-I | FT-II |
| --- | --- | --- | --- |
| Full-parameter fine-tuning | 0.26 | 0.74 | 0.28 |
| Freezing-layer fine-tuning | 0.26 | 0.55 | 0.33 |
| Adapter tuning | 0.26 | 0.65 | 0.52 |
| LoRA | 0.26 | 0.70 | 0.56 |
| LoRAMoE | 0.26 | 0.83 | 0.71 |

### S6 Structural characteristics of screened peptide binders

**Table S13.** Structural characteristics of the remained 157 peptide binders after static structure screening.

| Structural characteristics | Helix number | Helix angle | Helix distance | Interface residue | Targeting atom | Miss-targeting atom | Hydrogen bond |
| --- | --- | --- | --- | --- | --- | --- | --- |
| Value | 3.0000 | 15.7331 | 10.4501 | 63.9398 | 442.0723 | 17.9638 | 36.0241 |

### S7 Molecular dynamics simulation results

Both radius of gyration (Rg) and root mean square deviation (RMSD) curves in Fig.S2 970 indicate that all the molecular dynamics simulations of the studied complexes reach 971 their dynamic equilibration during the last 50 ns. The binding energies under two 972 different models, solvation free energies (G solvation), and specific G solvation are 973 shown in Table.S14, Table.S15, Table.S16, and Table.S17, respectively.

**Fig. S2.**
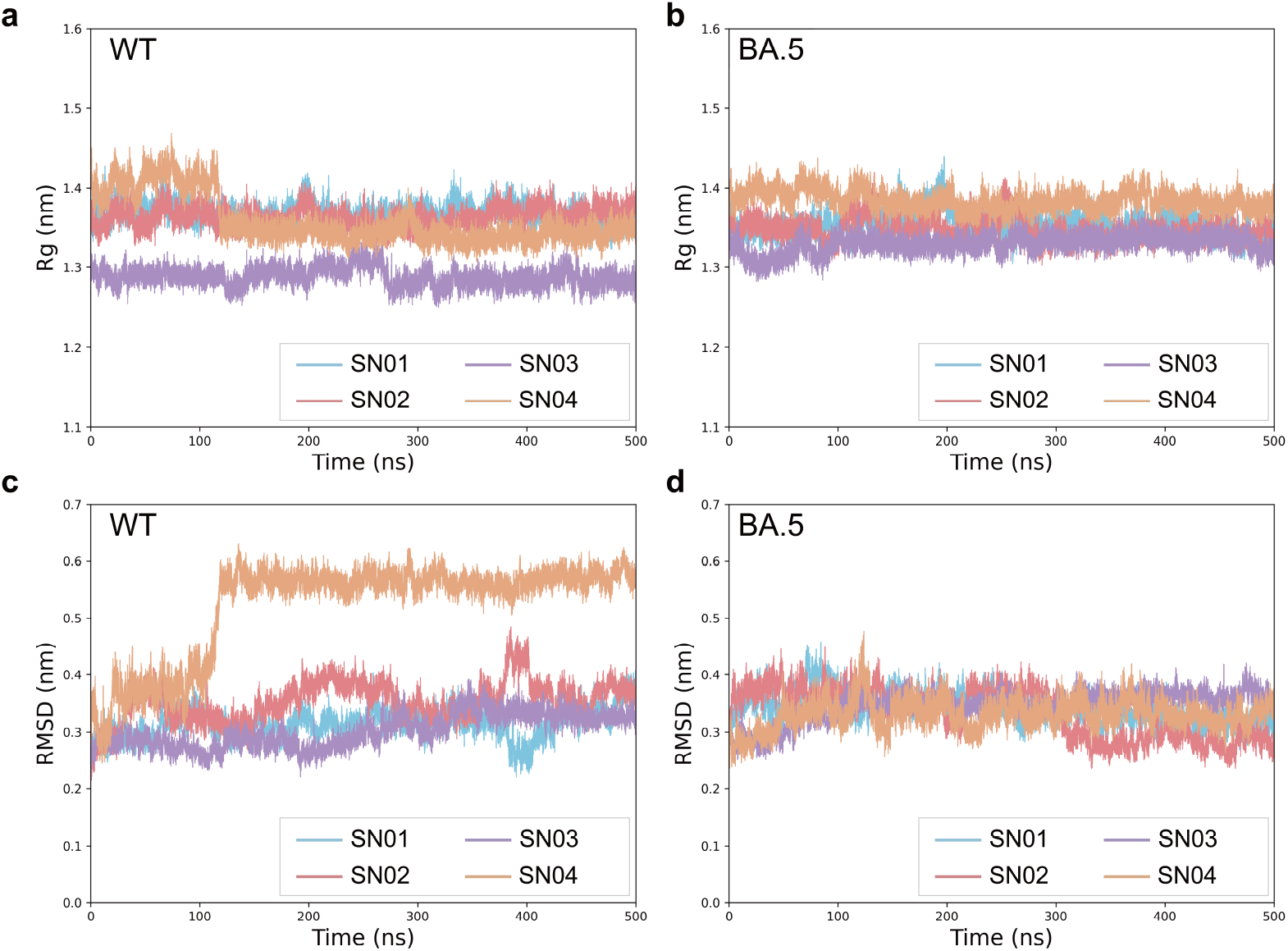
The radius of gyration (Rg) and root mean square deviation (RMSD) curves during molecular dynamics simulations. “WT” denotes the Rg (a) and RMSD (c) curves of the four most promising candidates with wild-type RBD as the target protein. “BA.5” denotes the Rg (b) and RMSD (d) curves of the four most promising candidates with Omicron BA.5 RBD as the target protein.

**Table S14.** Binding energies of the four most promising candidates to the target protein under SASA model.

| Complex | Binding energy |
| --- | --- |
| SN01&Wild-type RBD | -936.893±4.259 |
| SN01&Omicron BA.5 RBD | -1884.778±4.446 |
| SN02&Wild-type RBD | -714.377±4.915 |
| SN02&Omicron BA.5 RBD | -1277.094±4.114 |
| SN03&Wild-type RBD | -394.496±4.346 |
| SN03&Omicron BA.5 RBD | -756.901±6.224 |
| SN04&Wild-type RBD | -343.679±4.357 |
| SN04&Omicron BA.5 RBD | -854.127±4.943 |

**Table S15.**
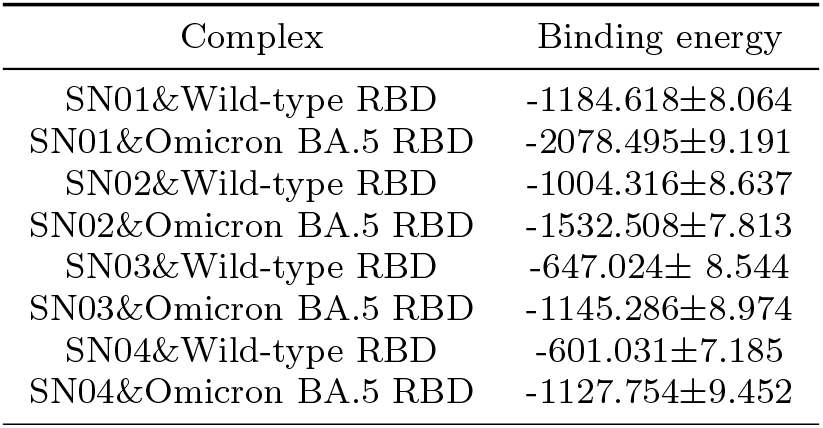
Binding energies of the four most promising candidates to the target protein under SAV model.

| Complex | Binding energy |
| --- | --- |
| SN01&Wild-type RBD | -1184.618±8.064 |
| SN01&Omicron BA.5 RBD | -2078.495±9.191 |
| SN02&Wild-type RBD | -1004.316±8.637 |
| SN02&Omicron BA.5 RBD | -1532.508±7.813 |
| SN03&Wild-type RBD | -647.024± 8.544 |
| SN03&Omicron BA.5 RBD | -1145.286±8.974 |
| SN04&Wild-type RBD | -601.031±7.185 |
| SN04&Omicron BA.5 RBD | -1127.754±9.452 |

**Table S16.** Solvation free energies (G solvation) of the four most promising candidates and two RBDs.

| Protein | G_solvation | SASA area |
| --- | --- | --- |
| Wild-type RBD | -5214.28236±179.15037 | 9884.544011±163.0892788 |
| Omicron BA.5 RBD | -6247.002615±206.734815 | 9845.216499±28.16386369 |
| SN01 | -10575.64074±211.37592 | 5821.713213±125.9938377 |
| SN02 | -6786.61242±188.63901 | 5702.487001±103.4844729 |
| SN03 | -5122.94601±160.94526 | 5064.135575±94.74562086 |
| SN04 | -5979.08808±170.52903 | 6086.440242±5.331034294 |

**Table S17.** Specific G solvation of the four most promising candidates and two RBDs.

| Protein | Specific G_solvation |
| --- | --- |
| Wild-type RBD | -0.527518756±0.000111131 |
| Omicron BA.5 RBD | -0.634521609±0.000745583 |
| SN01 | -1.81658566±0.000288174 |
| SN02 | -1.190114492±0.000319663 |
| SN03 | -1.011613124±0.000335439 |
| SN04 | -0.982362077±0.005255614 |

### S8 *In vitro* experimental validation results

**Table S18.**
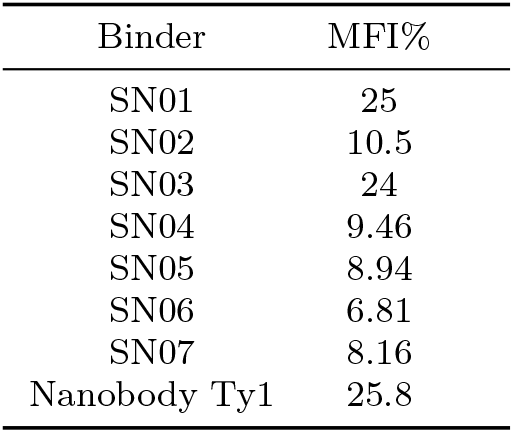
MFI% of screened 7 peptide binders.

**Table S19.**
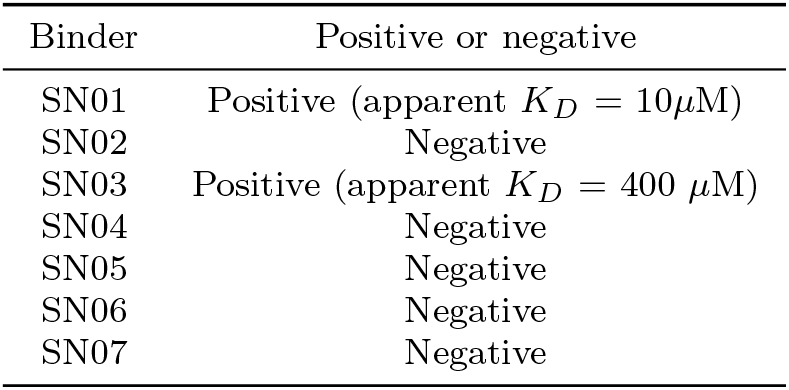
Binding affinity of screened 7 peptide binders targeting Omicron BA.5 RBD.

| Binder | Positive or negative |
| --- | --- |
| SN01 | Positive (apparent $K_D = 10\mu\text{M}$) |
| SN02 | Negative |
| SN03 | Positive (apparent $K_D = 400\mu\text{M}$) |
| SN04 | Negative |
| SN05 | Negative |
| SN06 | Negative |
| SN07 | Negative |

**Table S20.** Binding affinity of screened 7 peptide binders targeting wild-type RBD.

| Binder | Positive or negative |
| --- | --- |
| SN01 | Negative |
| SN02 | Negative |
| SN03 | Negative |
| SN04 | Negative |
| SN05 | Negative |
| SN06 | Negative |
| SN07 | Negative |
| Nanobody Ty1 | -(apparent $K_D = 17\text{ nM}$) |

## Footnotes

1 https://bfd.mmseqs.com

2 http://pfam.xfam.org/

3 https://www.uniprot.org/

